# Coordinating cell polarization and morphogenesis through mechanical feedback

**DOI:** 10.1101/2020.05.21.108076

**Authors:** Samhita P. Banavar, Michael Trogdon, Brian Drawert, Tau-Mu Yi, Linda R. Petzold, Otger Campàs

## Abstract

Many cellular processes require cell polarization to be maintained as the cell changes shape, grows or moves. Without feedback mechanisms relaying information about cell shape to the polarity molecular machinery, the coordination between cell polarization and morphogenesis, movement or growth would not be possible. Here we theoretically and computationally study the role of a genetically-encoded mechanical feedback (in the Cell Wall Integrity Pathway) as a potential coordination mechanism between cell morphogenesis and polarity during budding yeast mating projection growth. We developed a coarse-grained continuum description of the coupled dynamics of cell polarization and morphogenesis as well as 3D stochastic simulations of the molecular polarization machinery in the evolving cell shape. Both theoretical approaches show that in the absence of mechanical feedback (or in the presence of weak feedback), cell polarity cannot be maintained at the projection tip during growth, with the polarization cap wandering off the projection tip, arresting morphogenesis. In contrast, for mechanical feedback strengths above a threshold, cells can robustly maintain cell polarization at the tip and simultaneously sustain mating projection growth. These results indicate that the mechanical feedback encoded in the Cell Wall Integrity pathway can provide important positional information to the molecular machinery in the cell, thereby enabling the coordination of cell polarization and morphogenesis.

**Author summary:** Cell migration, morphogenesis and secretion are among the vast number of cellular processes that require cells to define a preferred spatial direction to perform essential tasks. This is achieved by setting an intracellular molecular gradient that polarizes the cell. While the molecular players involved in cell polarization and some of the mechanisms that cells use to establish such molecular gradients are known, it remains unclear how cells maintain polarization as they dramatically change shape during morphogenesis, migration, etc. Here we identify a potential feedback control mechanism, encoded genetically in cells, that provides the molecular polarization machinery with the necessary information about cell geometry to maintain cell polarization during cell shape changes.

## Introduction

Cell polarization is essential to a large number of cellular processes. From cell migration in animal cells to the growth of walled cells from plants and fungi, cells need to spatially polarize to perform key functions [1–7]. In many cases, an external cue, such as a molecular gradient, triggers the molecular polarization of the cell. Once polarity is established, cells need to maintain polarization at specific spatial locations on the cell surface to properly change shape or move in the right direction, suggesting the existence of a coordination mechanism between the molecular polarization machinery and the physical/geometrical changes of the cell. In animal cells, it has been proposed that membrane tension could help such coordination [1, 8–10], but no specific coordination mechanisms are known [7, 11, 12]. More generally, very little is known about the mechanisms that provide the molecular polarization machinery with the necessary information about cell geometry and how such polarization machinery coordinates with cell shape changes.

During polarized (tip) growth in walled cells, a process occurring in many different species ranging from plants and fungi to bacteria, cells must maintain polarity at the growing apex [13–17]. An example of this process is mating projection growth in budding yeast, *Saccharomyces cerevisiae* (Fig. 1a). During mating, *a*-cells and *α*-cells secrete pheromones, *a* factor and *α* factor, respectively, to attract the opposite cell type [18–20]. G protein coupled receptors, Ste2, on the cell surface bind pheromone molecules, triggering a cascade of reactions ultimately leading to the spatial localization of the polarity master regulator Cdc42 (Fig. 1b). In turn, Cdc42 recruits Bni1, a formin, which initiates the nucleation of actin cables from the polarization cap (Fig. 1c). These actin cables focus transport to the polarization cap, bringing more Cdc42 as well as the enzymes to synthesize and remodel the cell wall (Fig. 1d). This process induces the localized growth of the mating projection in the direction of maximal pheromone gradient.

**Fig 1.**
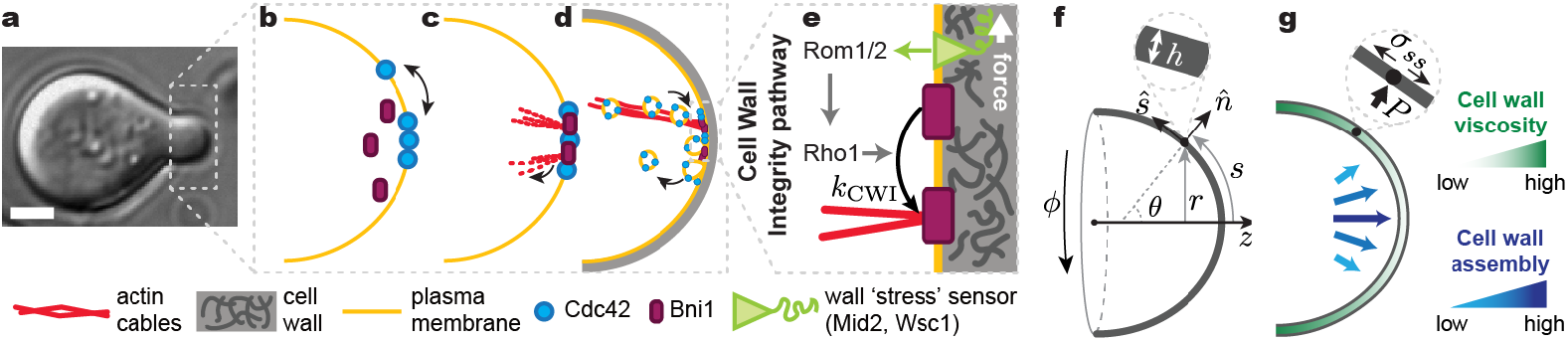
Schematic representation of relevant cell polarization events during budding yeast mating projection growth. **a**, Transmitted light image of a *S. cerevisiae* cell growing a mating projection in the presence of *α*-factor. Scale bar, 2*μ*m. **b-e**, Sketch of molecular events leading to the polarization of the cell during the growth of the mating projection. **f**, Geometrical representation of the system and definition of the relevant variables. **g**, Sketch depicting the increasing cell wall viscosity and decreasing cell wall assembly away from the apex. The inset depicts local normal force balance at the cell wall. All variables are defined in the main text.

While cell polarization and subsequent mating projection growth do occur in the presence of pheromone gradients, a graded cue is not necessary. If the distribution of *α*-factor is uniform (no gradient), the molecular polarization machinery has been shown to be able to spontaneously break the symmetry and polarize the cell [21], albeit in a random spatial direction. Upon polarization, a mating projection emerges from the polarization cap, which is maintained at the tip of the growing projection as it extends [22]. In the absence of a pheromone gradient to externally guide cell polarization and cell shape changes, the observation of sustained mating projection growth suggests the existence of a cell autonomous coordination mechanism between cell polarity and cell shape. However, it is unclear what, if any, cell autonomous coordination mechanism is at play during mating projection growth.

Many existing models of cell polarization in budding yeast are able to reproduce the spontaneous symmetry breaking and establishment of a polarization cap in a static, spherical cell geometry [21, 23–29]. However, using 3D simulations of the polarization machinery in non-spherical geometries, we have recently shown that these models cannot explain the maintenance of cell polarization at the tip of mating projections [30]. A polarization cap initially localized at the tip of the mating projection, quickly moves away from the tip and localizes out of the mating projection, thereby precluding mating projection growth. In these simulations, the polarization machinery is coupled to the cell geometry because the molecular gradients established inside the cell depend on the cell’s shape and, in turn, these geometry-dependent gradients affect the reaction rates and spatial distribution of polarization molecules at the cell surface. However, this polarization-cell geometry coupling does not help maintain cell polarization at the tip of the mating projection, but rather drives it away from that point [30]. A different coordination mechanism between cell shape and polarization must thus exist to maintain polarization at the projection tip and sustain mating projection growth.

Cell shape and cell wall mechanics are directly related to each other and, therefore, the mechanics of the cell wall contains information about the cell shape. We have recently shown that the sustained growth of a mating projection requires a mechanical feedback between the cell wall synthesis and cell wall mechanics [31]. This mechanical feedback is genetically encoded in the Cell Wall Integrity (CWI) pathway and involves cell wall “stress” sensors (Wsc1, Wsc2 and Mid2) activating the rho GTPase Rho1 (Fig. 1e), which, in turn, activates cell wall synthesis (Fks1/2), thereby creating a feedback loop between cell wall mechanics (cell shape) and cell wall assembly. In addition, Rho1 activation through the CWI pathway also promotes actin cable formation via the local activation of the formin Bni1 (Fig. 1e). Since Cdc42 is transported on actin cables by vesicles, it is possible that the same mechanical feedback shown to stabilize mating projection growth is simultaneously used to maintain cell polarity at the mating projection tip, effectively providing the molecular polarization machinery with the necessary positional information.

Unlike previous theoretical descriptions, which either considered the dynamics of polarization in fixed geometries [21, 23–29] or the mechanics of cell morphogenesis without accounting for polarization [31–34], here we couple the dynamics of cell polarization and cell wall mechanics during mating projection growth. Using both a coarse-grained theoretical description and 3D stochastic simulations of cell polarization coupled to the mechanics of cell morphogenesis, we show that the mechanical feedback encoded in the CWI pathway can, by itself, coordinate the dynamics of cell polarization and morphogenesis, maintaining the polarization cap at the tip of the mating projection and sustaining mating projection growth.

## Materials and methods

### Numerical integration of coarse-grained continuum equations

The system of equations was scaled and written in a manner such that *r*, *h*, *ρ*_C_ and *ρ*_A_ were described by time-dependant equations, and *u*, *θ*, *κ*_s_ by differential equations in *s*. The latter equations were solved by the method of lines; *s* was discretized and the *s*-derivatives were written as a differential matrix using fourth order central difference and one sided finite differences at the boundary. The resulting system becomes a differential algebraic system (DAE), which was solved using Sundials [35], a suite of nonlinear and DAE solvers. Steady state solutions were obtained by ensuring that all time derivatives of scaled variables were below 10^−3^.

### Numerical integration of coupled reaction diffusion simulations

The mechanics equations were coupled to the reaction diffusion equations. Because of the separation of timescales of the physical growth governed by the mechanics equations and the biochemical reactions governed by the stochastic reaction diffusion equations, we were able to use a split-operator approach to solve the coupled dynamics of cell wall expansion and the molecular polarization. The mechanics equations were solved using the same method presented above for the integration of the coarse-grained continuum equations. The stochastic reaction diffusion equations were built and solved using PyURDME and MOLNs [36]. Computational 3D meshes for each geometry consisted of a discretization of both the cytoplasm and the membrane (surface defined by cell shape), allowing for diffusion both in the cytoplasm and on the membrane as required by the models used in this study. All reactions took place in voxels on the membrane for each geometry. These meshes were generated using Gmsh [37].

## Results

To study the relation between the dynamics of cell polarization and cell morphogenesis during budding yeast mating projection growth, we combine different methodologies. First, we describe the mechanics of cell wall expansion that governs cell shape changes and, in particular, the extension of a mating projection. We then describe the dynamics of cell polarization using two approaches: (1) a continuum minimal model for the coupling between polarization and mechanics and, (2) 3D stochastic simulations of the molecular polarization machinery in the growing cell. In both cases, we couple the dynamics of polarization to the mechanics of the cell wall through the feedback encoded in the CWI pathway and study whether cell polarization can be maintained at the tip of the extending mating projection and whether mating projection growth can be sustained by the coupled dynamics.

### Geometry and mechanics of cell wall expansion during mating projection growth

As in any walled organism, the budding yeast cell is surrounded by a thin cell wall (~ 100 nm [38]), much smaller than the characteristic cell size or diameter of the mating projection (~ 1 *μ*m [39]). The shape of the cell is defined by the location of the cell wall and, during mating projection growth, it can be approximated to be axisymmetric (Fig. 1a,f). The extension of the mating projection can thus be described as the expansion of an axisymmetric thin shell caused by the large cell’s internal turgor pressure, *P* (Fig. 1f,g). It is convenient to parametrize the geometry of the cell (cell wall) by the arclength *s* from the projection apex and azimuthal angle *ϕ* (Fig. 1f). The shape of the growing projection is characterized by its local radius of curvature, *r*(*s*, *t*), and the two principal curvatures *κ_s_* = *∂θ*/*∂s* and *κ_ϕ_* = sin *θ*/*r*, respectively, where *θ*(*s*, *t*) is the angle between the local outward normal and the axis of growth (Fig. 1f). The coordinates (*r*, *ϕ*, *z*) are standard cylindrical coordinates, and the angle *θ* and arclength *s* provide measures of changes in the normal and tangential directions of the surface, 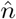 and 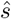 respectively [32, 40] (Fig. 1f).

The dynamics of cell wall expansion during growth are governed by mass and momentum conservation, as explained in previous works on tip growth [31, 32] and on the expansion of thin viscous shells [40]. Since inertia is irrelevant in this system, momentum conservation reduces to local force balance on the cell wall, which yields

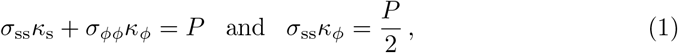

where *σ*_ss_(*s*, *t*) and *σ_ϕϕ_*(*s*, *t*) are the tensions along *s* and *ϕ* directions in the wall (Fig. 1f). The rheology (mechanical properties) of the cell wall control its expansion in response to the tensions in the wall. While the yeast cell wall in known to behave elastically at short time scales (seconds [38]), plastic behavior manifested by irreversible cell wall expansion occurs at the growing tip on the timescales of mating projection growth (minutes [41]). Enzymes that locally degrade the cell wall (glucanases) are brought to the polarization cap by exocytic vesicles moving along the actin cables emanating from it [42] (Fig. 1d). Upon release to the adjacent cell wall, a higher concentration of glucanases degrade more the cell wall adjacent to the polarization cap, creating a gradient of cell wall degradation away from it. This is consistent with the observation of a higher concentration of cell wall degrading enzymes (glucanases) near the apex of the mating projection [43]. Therefore, at the timescales of growth, the cell wall can thus be approximated by a viscous fluid shell with inhomogeneous viscosity, *μ*(*s*, *t*), minimal at the polarization cap and increasing away from it (Fig. 1g). In this case, the local tangential velocity *u*(*s*, *t*) along the arclength 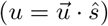 of a cell wall with constant density *ρ*_w_, or equivalently, its strain (expansion) rates 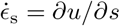 and 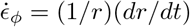, are related to the tensions in the cell wall by [32, 40]

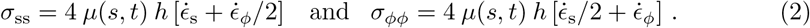

Beyond the expansion of the cell wall in response of the tensions in it, new cell wall needs to be assembled as the mating projection grows. The cell wall is assembled via Fks1/2 synthases at the plasma membrane that extrude 1, 3 – *β* glucans to the preexisting wall [44, 45]. Since Fks1/2 synthases are also brought to the plasma membrane along actin cables and inserted in it during the exocytosis process, their concentration is high at the polarization cap and decreases away from it, generating a graded distribution G(*s*, *t*) of the cell wall assembly rate that decreases away from the polarization cap (Fig. 1g). Mass conservation of the cell wall material during mating projection growth yields [31]

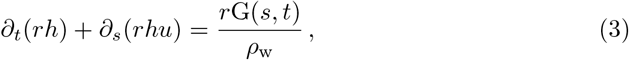

where *h*(*s*, *t*) is the cell wall thickness (Fig. 1e) and *ρ*_w_ is the constant cell wall density.

The functions *μ*(*s*, *t*) and G(*s*, *t*) depend directly on transport along actin cables and exocytosis, which are controlled by the polarization machinery. We will relate these functions to the dynamics of polarization below.

### Dynamics of cell polarization: minimal coarse-grained description

In order to form a mating projection, the cell must first polarize to specify the site of cell wall growth and expansion. Many molecular players are involved in the cell polarization process [46]. In our 3D stochastic simulations (see below), we will consider the coupled dynamics of some of the key molecular players in the polarization process. However, since the aim of the coarse-grained description is to study solely the coupling between the dynamics of polarization and the mechanics of cell wall expansion during mating projection growth, we neglect much of the complexity of the polarization molecular machinery in this minimal description. Since the formin Bni1 is directly activated through the CWI pathway, leading to the assembly of actin cables and further recruitment of the master polarity regulator Cdc42 via actin-mediated transport, we consider solely the effective, coupled dynamics of Cdc42 and Bni1 on the curved geometry of the cell.

Cdc42 can reach the plasma membrane either through direct binding from a cytoplasmic pool [47, 48] or through actin cable-mediated transport, carried by secretory vesicles [48]. Once at the membrane, Cdc42 diffuses with diffusion coefficient *D* and also unbinds from the membrane at a rate *k_D_*. The Cdc42 population at the plasma membrane acts as a central regulator of many polarization factors [47] and, in particular, it recruits Bni1, a formin that drives the nucleation of actin cables [49] (Fig. 1c). As mentioned above, these actin cables bring many different factors that are essential for cell wall remodeling and growth, as well ad Cdc42 itself, creating a positive feedback loop. Focusing on this feedback between Cdc42 and Bni1, we can write the dynamics of Cdc42 on the plasma membrane of the mating projection as

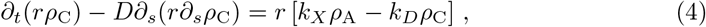

where *ρ_C_* is the Cdc42 concentration on the plasma membrane, *k_X_* is the rate at which Cdc42 is added to the plasma membrane through transport along actin filaments, and *ρ*_A_ is the local surface density of actin cables. Since actin cables are nucleated by active Bni, we assume the local density of actin cables emanating from the membrane to be proportional to the local active Bni1 concentration, *ρ*_B,a_, at the plasma membrane, namely *ρ*_A_ ~ *ρ*_B,a_. In this minimal description, we assume a uniform pool of inactive Bni1 on the plasma membrane with concentration *ρ*_B,i_ = *ρ*_0_ and write the dynamics of active Bni1 as

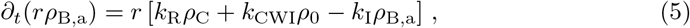

where *k_R_* is the rate at with Cdc42 recruits (and activates) Bni1, *k*_I_ is the inactivation rate of Bni1 and *k*_CWI_ is rate of activation of Bni1 through the CWI pathway (independently of Cdc42) [50]. Activation of the CWI pathway depends on the local mechanical state of the cell wall, leading to a direct coupling between the dynamics of Bni1 and the mechanics of the cell wall. Since CWI activation correlates with locations of cell wall expansion, we write the local activation rate of Bni1 via the CWI as being proportional to the local cell wall expansion (strain) rates [31], namely

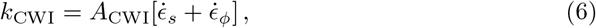

where *A*_CWI_ is a dimensionless constant measuring the strength of the mechanical feedback.

In order to fully connect the dynamics of Cdc42 and Bni1 to the mechanics of cell wall expansion during morphogenesis, it is necessary to relate them to the cell wall viscosity *μ*(*s*, *t*) and the rate of new cell wall assembly G(*s*, *t*). Since cell wall synthases Fks1/2 are carried to the plasma membrane through secretory vesicles along actin cables, we write the rate of new cell wall assembly G(*s*, *t*) to be proportional to the local concentration of actin cables, namely G(*s*, *t*) = *k_s_ ρ*_A_, with *k_s_* being the Fks1/2 rate of new wall assembly. Similarly, the spatial variations in cell wall viscosity reflect spatial changes in the concentration of glucanases, which are also transported to the cell surface through actin cables. Consequently, we assume the length scale of viscosity variations in the cell wall to be determined by spatial variations in actin cable density, namely 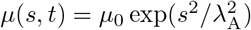, where λ_A_ is the length scale of the decay in actin cable density from the tip of the mating projection. This indicates that in regions with high concentration of actin cables, the cell wall viscosity is lower, promoting local cell wall expansion in that region.

Connecting the cell wall mechanics (Eqns. 1-3) to the dynamics of Cdc42 and Bni1 (Eqns. 4-5) as described above, we solved the coupled dynamics of polarity and cell wall expansion during cell morphogenesis. Scaling length and time with the length scale (*D*/*k_D_*)^1/2^ and time scale 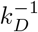 respectively, the stresses in the system with the turgor pressure *P* and the surface densities of Cdc42 and active Bni1 with *ρ*_0_, we obtain the relevant dimensionless parameters in the system (Table 1). The parameters *k*_I_/*k_D_*, *k*_R_/*k_D_*, *k_X_*/*k_D_* have either been measured or estimated and we use their known values (Supporting Information). To explore the role of a potential mechanical feedback between cell wall mechanics and polarization, we study the dynamics of the system upon changes in the feedback strength *A*_CWI_, which has not been directly measured.

**Table 1.**
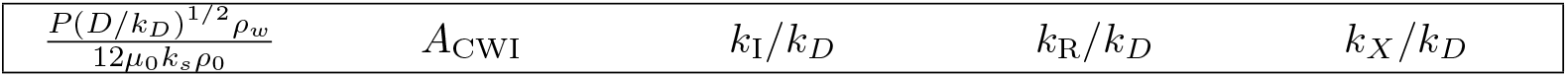
Relevant dimensionless parameters.

|  |  |  |  |  |
| --- | --- | --- | --- | --- |
| $\frac{P(D/k_D)^{1/2} \rho_w}{12\mu_0 k_s \rho_0}$ | $A_{CWI}$ | $k_I/k_D$ | $k_R/k_D$ | $k_X/k_D$ |

### Mechanical feedback can maintain cell polarization at mating projection tip

We first study the dynamics of polarization in the elongated shape of the growing projection, starting from the geometry of a mating projection. In the absence of mechanical feedback (*A*_CWI_ =0), an initially polarized cell loses its polarization within the typical timescales of molecular polarization processes, as can be seen from the loss in polarization in Cdc42 (Fig. 2a). Since the typical timescales of the molecular processes associated with polarization are shorter than the timescales of physical mating projection growth, cell polarization is lost before any cell shape changes occur. In contrast, if the strength of the mechanical feedback is large enough, cells maintain polarization at the tip of the growing mating projection. Even starting from a uniform distribution of Cdc42 in the mating projection (not polarized), the mere presence of the mechanical feedback polarizes the cell at the tip of the mating projection, enabling the sustained growth of the projection and the maintenance of polarization at the tip during growth (Fig. 2b). In between these two limiting regimes, there is a critical value of the mechanical feedback strength below which the cell is unable to maintain cell polarization (Fig. 2c), eventually leading to a uniform and very low (vanishing) concentration of Cdc42 at the plasma membrane. Above the critical value, stable solutions for the polarized state exist and the cell can maintain cell polarization at the tip of the mating projection during growth (Fig. 2b,c).

**Fig 2.**
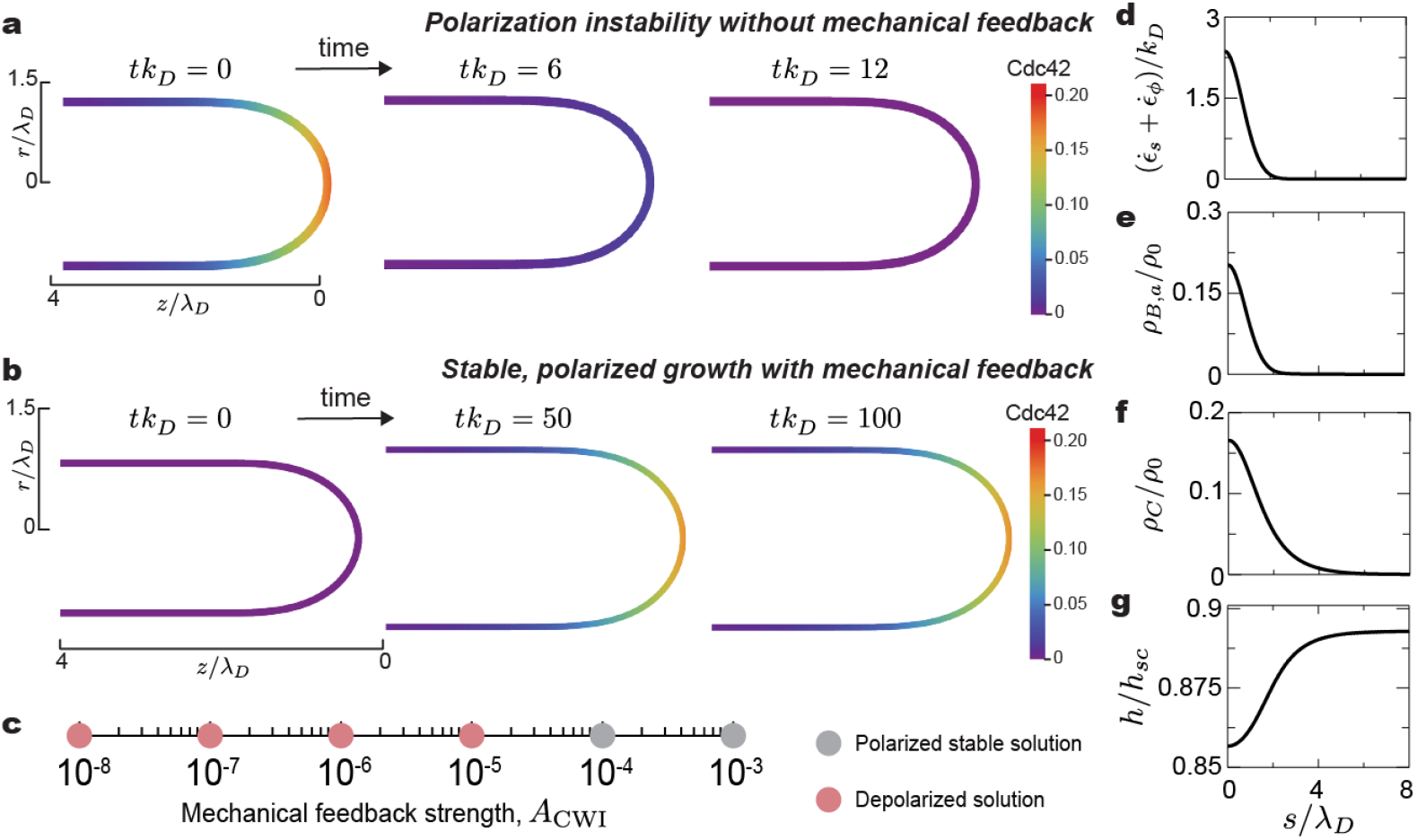
Maintenance of cell polarization at the mating projection tip via mechanical feedback. **a**, A cell initially polarized at the tip, as indicated by the distribution of Cdc42, loses polarization in the absence of mechanical feedback. The cell shape does not change over the short timescale of depolarization. **b**, An initially depolarized cell (uniform and low Cdc42 concentration) polarizes and grows steadily in the presence of mechanical mechanical feedback. **c**, Polarized or depolarized cell states for varying values of the mechanical feedback strength. Small feedback strengths lead to depolarized solutions (loss of Cdc42 localization at the tip). For feedback strengths above a threshold, cell polarization is stably maintained at the tip during cell growth (Polarized stable solution). **d-g**, Steady state spatial profiles for cell wall expansion rate 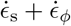 (**d**), active Bni1 density (**e**), Cdc42 density (**f**) and cell wall thickness *h* (**g**).

As expected for mating projection growth [31], the stable solutions show that the rate of cell surface (cell wall) expansion, namely 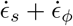, is maximal at the tip and decays away from it with the characteristic length scale λ_*D*_ of that defines the polarity cap (Fig. 2d), eventually vanishing far away from the tip, where the cell wall becomes solid-like (diverging viscosity; Fig. 1g). The presence of mechanical feedback, leads to the localized activation of Bni1 at the tip (Fig. 2e) that, in turn, generates a localized (polarized) density of actin cables emanating from the tip of the growing mating projection. This higher density of actin cables at the tip generates higher concentrations of Cdc42 in this region (Fig. 2f), maintaining cell polarization during cell growth. Finally, the cell wall thickness is largely uniform along the mating projection, albeit with a slight thinning at the tip (Fig. 2g), in agreement with experimental observations [51, 52].

This minimal description shows that the mechanical feedback between cell wall expansion and Bni1 dynamics encoded in the CWI pathway can maintain cell polarization at regions of the cell surface where cell wall expansion occurs. Since cell wall expansion and cell geometry are related to each other through mass and momentum conservation, this mechanical feedback effectively provides the cell polarization machinery with information on the cell shape (i.e., positional information).

### Stochastic simulations of cell polarization during mating projection growth

Because the timescales of cell wall expansion and growth (cell shape changes) are much longer than the timescales of the molecular reactions involved in cell polarization, we first studied the polarization dynamics by performing stochastic simulations of molecular polarization models in a fixed cell geometry. The shape of the cell with a growing mating projection was obtained from the mechanical description described above, following our previous work [31].

To perform 3D stochastic simulations of the polarization dynamics in the fixed cell geometry, we employed the previously described PyURDME software [36], which can simulate spatial stochastic dynamics on complex 3D and time-dependent geometries. In this stochastic model, we considered both the inactive (Cdc42-GDP) and active (Cdc42-GTP) states of Cdc42, the local density of actin cables on the membrane and actin monomers in the cytoplasm (Fig. 3a). Inactive Cdc42 in the cytoplasm can directly bind to the membrane at a rate *β*_2_ (*β*_2_ ≃ 0.28*μms*^−1^ [28]) or transported to it by actin cables (at rate *β*_1_), and has a membrane diffusion constant of 0.0053*μm*^2^*s*^−1^ [53]. It can also dissociate from the plasma membrane at a rate *β*_3_ (*β*_3_ ≃ 1*s*^−1^ [28]). At the plasma membrane, inactive Cdc42 can be activated and inactivated at rates 0.2*μm*^2^*s*^−1^ and 1*s*^−1^ [28], respectively. Local assembly of actin cables on the membrane is due to Bni1 activation, which can be caused either by active Cdc42 (*A*_on_ ≃ 0.197*μm*^3^*s*^−1^ [53]; Fig. 3a) or by Rho1 through the CWI pathway at a rate that depends on the local cell wall expansion, namely 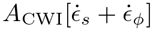 (Fig. 3d). Actin filaments disassemble at a rate *A*_off_ ≃ 2.70*s*^−1^ [53]. For a detailed description of the set of equations and parameters describing this model, please see the Supporting Information.

**Fig 3.**
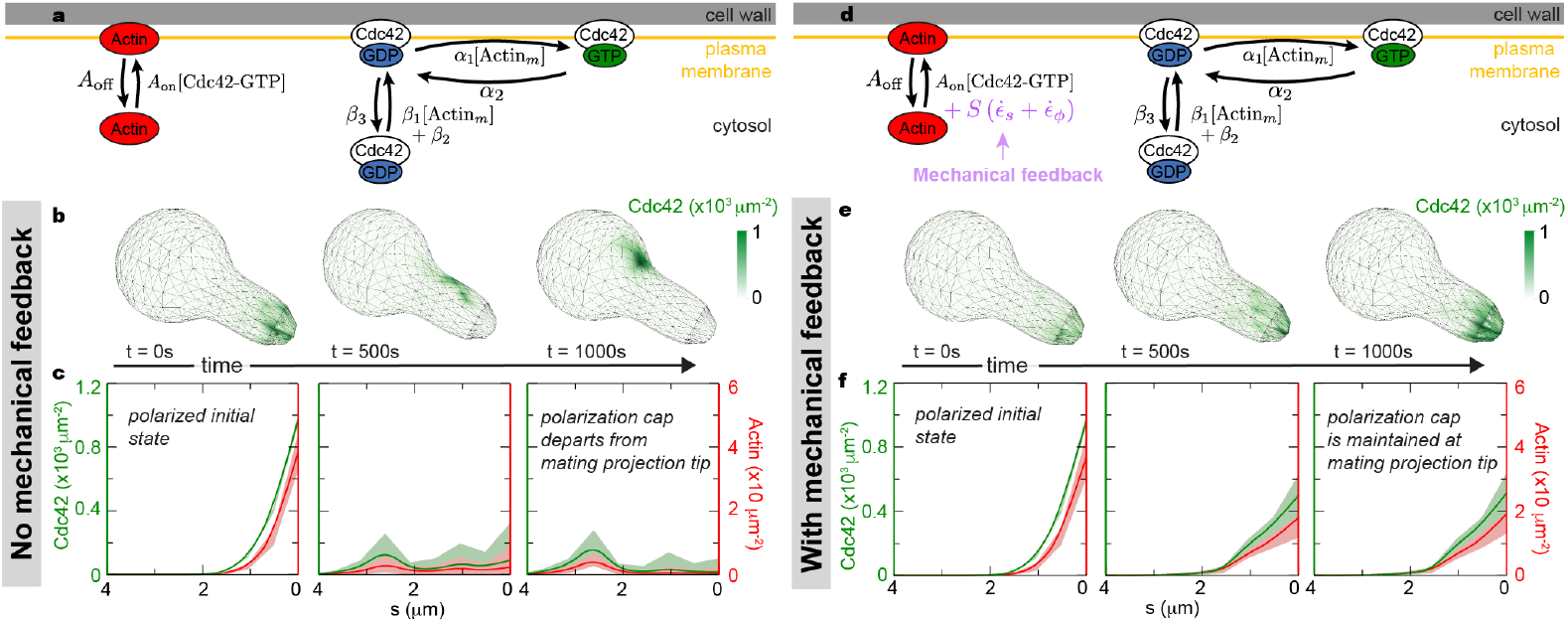
Role of mechanical feedback on the maintenance of the polarization cap in a fixed cellular geometry. **a**, Molecular events of the polarization machinery used in the 3D stochastic simulations, in the absence of mechanical feedback. The diagram focuses on reaction events and does not depict diffusion events. **b**, In the absence of mechanical feedback, a cell with the Cdc42 (green) polarization cap initially located at the tip, depolarizes over time, with the polarization cap delocalizing from the tip of the mating projection and moving towards the cell body. **c**, Averaged Cdc42 (green) and actin cable (red) densities along the cell arclength showing the loss of polarization at the tip of the mating projection over time in the absence of mechanical feedback. **d**, Molecular events of the polarization machinery used in the 3D stochastic simulations, in the presence of mechanical feedback. **e**, The polarization cap (high Cdc42 concentration) remains at the tip of the mating projection in the presence of mechanical feedback. **f**, Averaged Cdc42 and actin cable densities along the cell arclength showing that in the presence of mechanical feedback, the polarization cap remains at the tip. The 3D plots shown in **b** and **e** correspond to a single simulation realization, whereas the averaged profiless in **c** and **f** contain information of 100 realizations.

In the absence of mechanical feedback (*A*_CWI_ = 0), an initially polarized cell with high concentration of Cdc42 at the tip of the mating projection cannot maintain polarization at the tip (Fig. 3b-c). This polarization model can spontaneously break the symmetry and generate a polarization cap (Fig. 3b), as many other models of molecular cell polarization do in spherical geometries [21, 23–29]. However, as we have shown previously for various polarization models [30], the polarization cap moves away from the tip of the mating projection (Fig. 3b-c).

To study the effect of the mechanical feedback, we progressively increase its strength *A*_CWI_. For values of the mechanical feedback strength above a critical value, we observed that the polarization cap is maintained at the tip of the projection (Fig. 3e-f). While the shape of the cell is fixed (static) in these simulations, the values of the mechanical fields and, in particular, the cell wall expansion rate 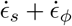 are those of the growing cell. Essentially, the molecular events associated with cell polarization are able to see a snapshot of the growing cell in these simulations. Since the cell wall expansion rate is localized at the tip of the mating projection [31] (Fig. 2d), the polarization machinery has a preferential location for actin cable assembly, bringing more Cdc42 to the tip and stabilizing the localization of the polarization cap in this region. These simulations show that the presence of the mechanical feedback on Bni1 activation, which is part of the CWI pathway, can maintain the polarization cap at the tip of the mating projection.

While the existence of a mechanical feedback can stabilize the polarization cap at the tip of a static mating projection (fixed geometry), it is unclear whether the polarization cap can also be maintained at the tip during mating projection growth. To address this point, we coupled the 3D stochastic simulations of cell polarization to the mechanics of the expanding cell wall. This involves performing the 3D stochastic simulations of the polarization machinery in a changing cell shape. To do so, we used an operator splitting methodology similar to that described in previous works [54], where the simulation domain can evolve over time in a predetermined manner (Methods). To specify how the simulation domain (cell shape) evolves over time, we used the description above of cell wall mechanics and expansion (Eqs. 1–3), as it governs the dynamics of cell shape changes. The operator splitting methodology takes advantage of the large separation of timescales between the fast molecular processes that polarize the cell and the slow physical cell wall expansion during cell growth.

Staring with a spherical geometry (sphere of radius *r* = 2*μm*) and random initial distribution for each chemical species (Fig. 4a), we simulate the polarization dynamics for timescales smaller than those leading to any relevant change in cell shape. The cell polarization machinery establishes a polarization cap within the spherical geometry (Fig. 4a), as observed experimentally [24] and recapitulated by many polarization models [21, 23–29]. This spontaneous symmetry breaking reduces the symmetry of the problem from a sphere to an axisymmetric geometry. Taking advantage of the axisymmetric geometry, we time and rotationally averaged the concentration of active Cdc42 and actin cables to obtain their distributions along the arclength *s* from the tip of the mating projection (Fig. 4b and Fig. 1f). Once these averaged distributions are known (Fig. 4b), they were used as input fields for the equations describing the mechanics of cell wall expansion. As described above in the coarse-grained description, we write the rate of new cell wall assembly G(*s*, *t*) to be proportional to the local concentration of actin cables, namely G(*s*, *t*) = *k_s_ ρ*_A_, and the cell wall viscosity 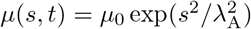, with λ_A_ being the length scale of the decay in actin cable density from the tip of the mating projection. In this specific simulations, the spatial profile of actin cable density *ρ*_A_(*s*) corresponds to the averaged distribution of actin on the membrane (Fig. 4b), and λ_A_ is the decay lengthscale obtained from the actin cable density profile (Fig. 4b). Knowing G(*s*) and *μ*(*s*) from the averaged actin cable density profile, we integrate the Eqs. 1–3 describing cell wall expansion from the initial spherical geometry and for a single simulation timestep. Since cell wall expansion is much slower than the molecular polarization dynamics, only a small (infinitesimal) cell shape change occurs. We then continue the 3D stochastic simulations of polarization in this new cell geometry (Methods), leading to small changes in the spatial profile of Cdc42 and actin cables. The new averaged spatial profile of actin cable density is then used to obtain the new G(*s*) and *μ*(*s*) and we solve again the mechanics of cell wall expansion. Iteration of this process allows us to simulate the stochastic dynamics of polarization in the evolving cell shape.

**Fig 4.**
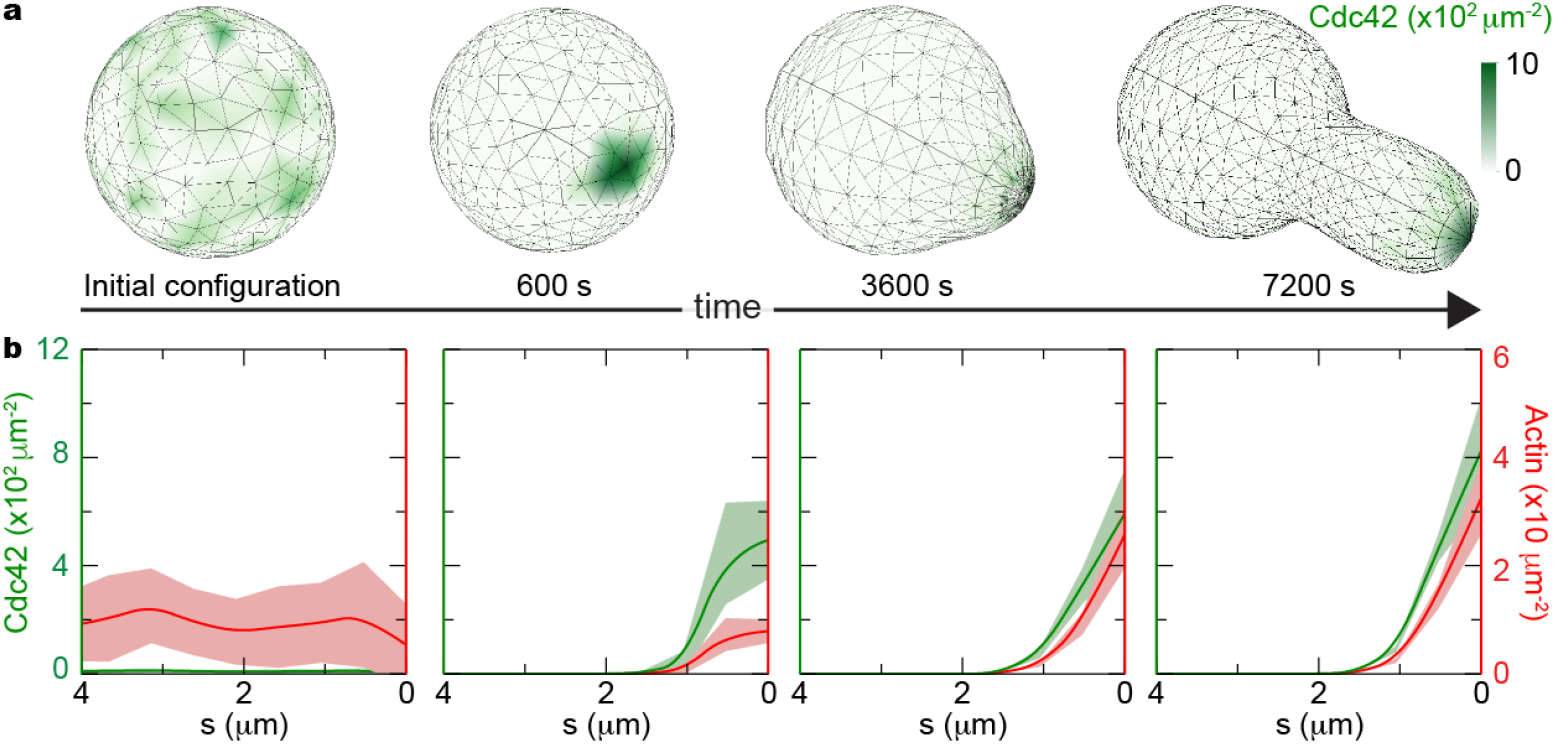
Coupled dynamics of cell polarization and cell wall expansion during mating projection growth in the presence of mechanical feedback. **a**, Time evolution of the 3D cell shape and Cdc42 distribution on the cell surface for a single realization of the stochastic simulation. Starting from a uniform density of Cdc42 (green) in a spherical geometry, the cell polarizes and grows a mating projection, maintaining the polarization cap at the tip of the growing mating projection. **b**, Time evolution of the averaged surface Cdc42 (green) and actin cable (red) densities along the cell arclength the spontaneous polarization of the cell from a depolarized state (uniform Cdc42 concentration) and the maintenance of cell polarization at the tip of the mating projection during growth, as indicated by the maximal values of Cdc42 and actin cables densities at the tip. These profiles correspond to averages over 100 realizations.

To understand whether the mechanical feedback encoded in the CWI pathway could also stabilize the polarization cap during mating projection growth, we simulated their coupled dynamics as described above. For values of the mechanical feedback strength similar to those that stabilized the polarization cap in a fixed geometry, the polarization cap is maintained stable at the tip of the mating projection during growth (Fig. 4a). The profiles of both Cdc42 and actin cables are polarized at the tip (Fig. 4b), displaying a higher concentration in this region and decaying away from it. In the absence of mechanical feedback, no mating projection grows from the initial spherical cell because it is not possible to maintain the polarization cap stably located at the point where the mating projection starts to form. While the averaged profiles of both Cdc42 and actin cable density show very clear polarization at the projection tip (Fig. 4b), snapshots of a single simulation show more fluctuating profiles (Fig. 4a), as expected from stochastic simulations.

## Discussion

We theoretically explored how a mechanical feedback between the mechanics of cell wall expansion and the dynamics of the polarization machinery can stabilize and maintain cell polarization at the tip of a growing mating projection. Using both a coarse-grained description and full 3D stochastic simulations of polarization, our results show that a mechanical feedback encoded in the CWI pathway, which activates the formin Bni1 through mechanical input from the cell wall, can maintain the polarization cap at the tip of a growing mating projection. Our description indicates that such genetically-encoded mechanical feedback can provide important positional information to the molecular machinery in the cell, enabling the coordination of cell polarization and morphogenesis.

The minimal coarse-grained description showed the existence of a threshold in feedback strength, above which a stable polarization cap can be maintained at the tip. This result is similar to that obtained using 3D stochastic simulations coupled to the mechanics of mating projection growth. However, while the stochastic simulations are able to reproduce both the spontaneous symmetry breaking (spontaneous polarization of the cell) in the spherical geometry and also the maintenance of the polarization cap at the tip of the growing mating projection, the minimal coarse-grained description is unable to reproduce the spontaneous formation of a polarization cap. In the minimal description, even in the absence of a preformed polarization cap, the mechanical feedback can generate stably polarized solutions with a polarization cap at the tip. This is because cell polarization is established by the same feedback mechanism that maintains it at the tip. In contrast, the stochastic simulations show that the formation of the polarization cap is independent from the maintenance of the polarization cap at the tip of the projection. Despite these differences, both descriptions indicate that the mechanical feedback can maintain the polarization cap at the tip of a growing mating projection.

Previous models of cell polarization in budding yeast have focused solely on spherical geometries [23–27]. As the cell changes shape during mating projection growth, it is essential for the polarization cap to receive some positional information on the location of the growing tip or, alternatively, information about cell shape. Modeling the dynamics of cell polarization without any coupling to variables able to provide positional or geometrical information, as previously done, can properly reproduce the spontaneous symmetry breaking [24] (Fig 5a), but not the coordination between the polarization cap and cell shape (Fig 5b). While curvature-sensing proteins could potentially localize at the tip of mating projections and provide a scaffold to maintain polarity at the growing tip [55], no curvature-sensing proteins have been found in mating projections yet. In budding yeast, curvature sensing proteins are involved in the fusion of mating projections [56], but these have a preference for zero curvature surfaces and control proper cell fusion. The formation of actin cables could also help restrict the motion of the polarity cap on the cell surface, as previously suggested for a spherical cell [27], but it does not provide a clear feedback to maintain the polarity at specific locations on the cell surface. Cell-sized actin cables could have their configurations restricted by cell shape, leading to an effective coupling between cell geometry and polarity. However, such a coupling would not provide information on the state of the cell wall, which is very important for cell shape changes. Our description provides a mechanism to explain the maintenance of cell polarization at specific locations of the cell surface as the cell changes shape. We propose that the genetically-encoded mechanical feedback through the CWI pathway informs the polarization machinery of the location of the cell surface where the cell wall is expanding, thereby providing the necessary positional information to maintain polarity at the growing mating projection tip (Fig 5b).

**Fig 5.**
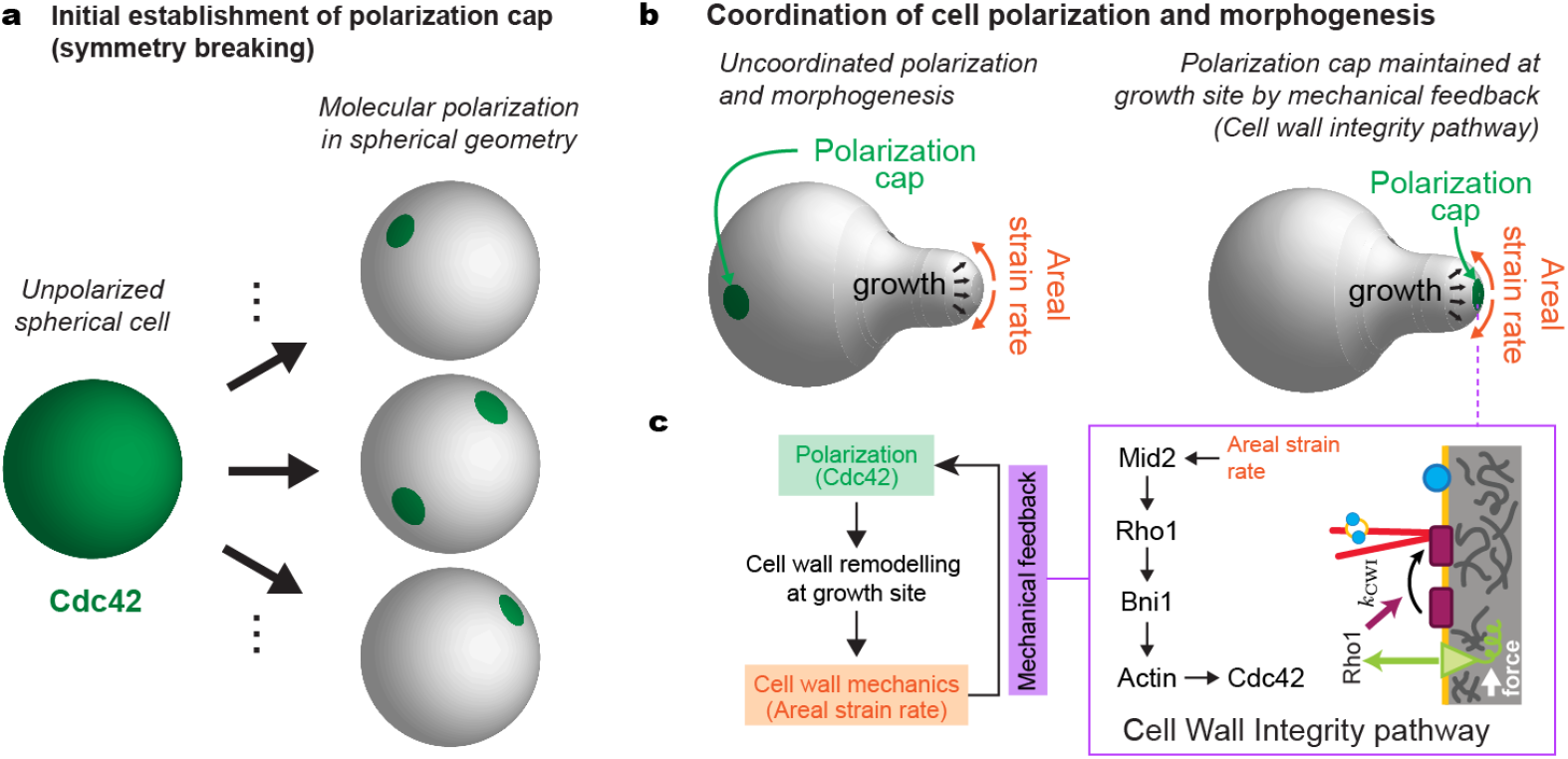
Coordination of cell polarization and mating projection growth via mechanical feedback. **a**, Unpolarized, spherical budding yeast cells can spontaneously break the symmetry and establish a polarization cap that is randomly located on the cell’s surface (all locations on the cell surface are equally probable) or even create multiple polarization caps. **b**, In the absence of any feedback between cell shape/mechanics and cell polarity, the polarity cap lacks positional information and cannot be maintained at the tip. In the presence of the mechanical feedback provided by the CWI pathway (see details of the feedback events in the box), the polarization cap is maintained at the tip of the growing mating projection. In this case, the polarization cap receives positional information through the mechanical feedback in the CWI pathway, which differentiates different positions on the cell surface associated with different levels of cell wall expansion.

Beyond budding yeast, many other organisms, including other fungi, plants and bacteria, undergo polarized cell wall growth [57–60]. The molecular control of cell wall remodeling and morphogenesis differs across species [61], and it is therefore possible that different mechanisms coordinate cell polarity and morphogenesis in other species. However, the mechanical feedback described herein for mating projection growth in budding yeast may also be present in closely related species, especially fungi like fission yeast [62, 63]. Indeed, fission yeast exhibits polarized growth and it has been recently shown that stable polarization caps are destabilized by the arrest of growth [12], suggesting the existence of a similar mechanical feedback. Given that in animal cells the actin cortex plays a similar role to the cell wall in walled cells, it is possible that a mechanical feedback between cortex expansion and polarity coordinates the molecular machinery with cell shape, thereby providing important positional information to intracellular processes. In early *C. elegans* development, there is a direct physical coupling between actin flows and the polarization machinery [11, 64–67], but it is unclear if a genetically-encoded mechanical feedback exists. Finally, membrane tension has been suggested to globally coordinate intracellular processes. Specifically, membrane-tension gradients, which may strongly depend on the actin cortex dynamics (both in animal [1, 8–10] and walled cells [68]), could potentially provide similar mechanical feedback mechanisms to the one proposed herein.

We previously showed that the CWI pathway encoded a mechanical feedback that stabilized budding yeast mating projection growth through the coupling of cell wall expansion and assembly [31]. The same feedback mechanism was later found in fission yeast [62], suggesting that similar mechanical feedback control mechanisms may exist in multiple species. Altogether, our previous results [31] and those presented here suggest that the same mechanical feedback mechanism, encoded in the CWI pathway, informs the intracellular molecular machinery of cell shape changes, thereby maintaining polarization at the tip of the mating projection while simultaneously ensuring its sustained and robust growth. Since the CWI pathway also couples cell wall mechanics to other intracellular processes (like exocytosis, etc.) through Rho1, it is possible that this mechanical feedback simultaneously coordinates many intracellular processes with cell morphogenesis.

The need to coordinate molecular processes with cell shape changes and growth is a general problem beyond polarization. Identifying the molecular mechanisms enabling this coordination at different scales and in different organisms will substantially contribute to our understanding of cell dynamics.

## Supporting Information

**S1 File. Supporting notes.** Parameters values for the minimal coarse-grained description and details on the 3D stochastic simulations.

## Acknowledgments

We would like to thank all members of the Campàs, Petzold and Yi groups for insightful discussions. Research reported in this publication was supported by the National Institute of General Medical Sciences of the National Institutes of Health under award number R01GM113241. We acknowledge support from the NSF Center for Scientific Computing in the California NanoSystems Institute (CNSI) and the Materials Research Laboratory (MRL) at UCSB (NSF MRSEC DMR-1121053 and NSF CNS-0960316).

